# New insights into the involvement of residue D1/V185 in Photosystem II function in *Synechocystis 6803* and *Thermosynechococcus vestitus*

**DOI:** 10.1101/2024.10.01.616037

**Authors:** Alain Boussac, Julien Sellés, Miwa Sugiura, Robert L. Burnap

## Abstract

The effects of D1-V185T and D1-V185N mutations in Photosystem II (PSII) from *Thermosynechococcus vestitus* (formerly *T. elongatus*) and *Synechocystis* 6803, respectively, were studied using both EPR and optical kinetics. EPR spectroscopy reveals the presence of a mixture of a S_2_ state in a high spin configuration (S_2_^HS^) and in a low spin configuration (S_2_^LS^) in both mutants. In contrast to the S_2_^HS^ in the wild type, the S_2_^HS^ state in the D1-V185T mutant does not progress to the S_3_ state at 198 K. This inability is likely due to alterations in the protonation state and hydrogen-bonding network around the Mn_4_CaO_5_ cluster. Optical studies show that these mutations significantly affect proton release during the S_3_-to-S_0_ transition. While the initial fast proton release associated with Tyr_Z_^●^ formation remains unaffected within the resolution of our measurements, the second, and slower, proton release is delayed, suggesting that the mutations disrupt the hydrogen-bonding interactions necessary for efficient deprotonation of substrate water (O6). This disruption in proton transfer also correlates with slower water exchange in the S_3_ state, likely due to non-native hydrogen bonds introduced by the threonine or asparagine side chains at position 185. These findings point to a critical role of D1-V185 in regulating both proton transfer dynamics and water binding, underscoring a complex interplay between structural and functional changes in PSII.

## Introduction

Oxygenic photosynthesis in cyanobacteria, algae and higher plants converts solar energy into the chemical bonds of sugars and oxygen [1]. Photosystem II (PSII) begins this process by splitting water to obtain electrons in the form of reduced quinone, while generating a proton gradient and releasing O_2_. The mature PSII binds 35 chlorophylls *a* (Chl-*a*),two pheophytins (Phe-*a*), one membrane b-type cytochrome, one extrinsic c-type cytochrome (in cyanobacteria and red algae), one non-heme iron, two plastoquinones (Q_A_ and Q_B_), the Mn_4_CaO_5_ cluster, 2 Cl^−^, 12 carotenoids and 25 lipids [2,3]. In the cyanobacterium *Synechocystis sp.* PCC 6803 a 4^th^ extrinsic subunit, PsbQ, has also been found in addition to PsbV, PsbO and PsbU [4].

Among the 35 Chl-*a*, 31 are antenna Chls. When one is excited, the excitation energy is transferred to other chlorophylls until it reaches the key pigments in the photochemical reaction center: 4 Chl-*a* molecules, P_D1_, P_D2_, Chl_D1_, Chl_D2_ and 2 Phe-*a* molecules, Phe_D1_ and Phe_D2_. A few picoseconds after the formation of the excited *Chl_D1_, charge separation occurs, ultimately forming the Chl ^+^Phe ^−^ and then P ^+^Phe ^−^ radical pair states, *e.g.* [5,6]. Formation of the Chl ^+^Phe ^−^ radical pair was recently defined as a fast pathway (short-range charge-separation) in contrast with a slow pathway with P_D1_P_D2_ as the initial donor (long-range charge separation) that would result directly in the formation of the P_D1_^+^Phe_D1_^−^ radical pair [7].

After the charge separation, P_D1_^+^ oxidizes Tyr_Z_, the Tyr161 of the D1 polypeptide, which is then reduced by the Mn_4_CaO_5_ cluster. The electron on Phe ^−^ is then transferred to Q_A_, the primary quinone electron acceptor, and then to Q_B_, the second quinone electron acceptor. Whereas Q_A_ can be only singly reduced under normal conditions, Q_B_ accepts two electrons and two protons before leaving its binding site and being replaced by an oxidized Q_B_ molecule from the membrane plastoquinone pool, see for example [1,8–12] for a non-exhaustive list of recent reviews on PSII function. The Mn_4_CaO_5_ cluster, oxidized by the Tyr_Z_^•^ radical formed after each charge separation, cycles through five redox states denoted Sn, where n designates the number of stored oxidizing equivalents. The S_1_ state is stable in the dark, which makes S_1_ the preponderant state after the decay of the S_3_ and S_2_ states in the dark. When the S_4_ state is formed, after the 3^rd^ flash of light given on dark-adapted PSII, two water molecules bound to the cluster are oxidized, O_2_ is released and the S_0_-state is reformed, [13,14].

Thanks to the advent of serial femtosecond X-ray free electron laser crystallography and cryo-EM spectroscopy, structures of the Mn_4_CaO_5_ cluster have been resolved in the dark-adapted S_1_ state, the S_2_ and S_3_ states, in a redox state as close as possible to those expected taking into account the misses under flash illumination. In the S_1_ state, *i.e.* in the Mn^III^_2_Mn^IV^_2_ redox state, the Mn_4_CaO_5_ structure resembles a distorted chair, including a μ-oxo-bridged cuboidal Mn_3_O_4_Ca unit with a fourth Mn attached to this core structure *via* two μ-oxo bridges involving the two oxygen’s O4 and O5 [2]. Recently, important progress has been made in the resolution of the crystal structures in the S_2_, S_3_ [15–17] and S_0_ states [15,16]. Briefly, changes in the S_1_ to S_2_ transition more or less correspond to those expected for the valence change of the Mn4 from +III to +IV. Importantly, water molecules in the “O1” and “O4” channels, defined as such because they start from the O1 and O4 oxygens of the cluster, appeared localized slightly differently in S_2_ and S_1_. In contrast, in the S_2_ to S_3_ transition, major structural changes have been detected together with the insertion of a 6^th^ oxygen (named either O6 or Ox), possibly that one of W3 originally bound to the Ca site, bridging Mn1 and Ca. This oxygen is supposed to correspond to the second water substrate molecule and is close to the bridging oxygen O5 supposed to be the first water substrate molecule [16,17,19]. An important movement of the Glu189 residue would allow its carboxylate chain to make a hydrogen bond with the protonated form of the 6^th^ oxygen in S_3_ [16,17], see also [18] for a recent computational work.

EPR studies show the existence of multiple structural forms for each of the S_1_, S_2_, and S_3_ states. The S_1_ EPR signals seen with a parallel mode detection at *g* ∼ 4.8 and *g* ∼ 12 [20–22] were attributed to an orientational Jahn–Teller isomerism of the dangler Mn4 with the valence +III [23].

In S_2_, at least two EPR signals can be detected at helium temperatures. The first one has a low-spin *S* = 1/2 value, S_2_^LS^, characterized by a multiline signal (ML) made up of at least 20 lines separated by approximately 80 gauss, centered at *g* ∼ 2.0 and spread over roughly 1800 gauss [24–26]. The second configuration of S_2_ is a high-spin ground state, S_2_^HS^, with *S* ≥ 5/ 2. In plant PSII, S_2_^HS^ may exhibit either a derivative-like EPR signal centered at *g ∼* 4.1 [27,28] or more complex signals at higher *g* values [29,30].

An influential computational study [31] proposed that the *g* ∼ 4.1 signal originates from a Mn_4_CaO_5_ cluster with a closed cubane structure. However, this proposed closed cubane structure for this S_2_^HS^ configuration has never been observed by XFEL studies of PSII, either from plants or cyanobacteria. For cyanobacteria, this result is not surprising since the *g ∼*4.1 signal has not been detected until now. Instead, the S_2_^HS^ EPR signal has a derivative-like shape centered at *g* ∼ 4.8 [32,33]. Yet, it remains possible that the proposed configuration is a short-lived transient that the time-resolved studies have not captured due to limited post-flash sampling frequency.

Other computational studies [34–36] proposed a different model in which, starting from the S_2_^LS^ configuration, protonation of O4 leads to an *S* = 5/2 ground state when W1, one of the two water molecules, along with W2, bound to the dangling Mn4 in the Mn_4_CaO_5_ cluster, is present as an aquo ligand. The further deprotonation of W1 to form a hydroxo ligand would then give rise to an *S* = 7/2 ground state [34–36]. It was further proposed that the form *S* = 7/2 was required to progress to S_3_. Importantly, the S_2_ ^HS^ form detected at *g* ∼ 4.8 form, *i.e.* the *S* = 7/2 form, that corresponds to an open cubane structure in [34–36], is able to progress to S_3_ at low temperatures [32,33], whereas in plant PSII the g ∼ 4.1 state cannot [37].

In most centers, the S_3_-state exhibits a spin *S* = 3 ground state [38–40]. In this *S* = 3 configuration, the four Mn ions of the cluster have an Mn^IV^ formal oxidation state with an octahedral ligation sphere in an open cubane structure [40]. In this proposed magnetic configuration, the dangler Mn^IV^ (*S* = 3/2) is antiferromagnetically coupled to the open cubane motif Mn^IV^_3_ with a total spin value *S* = 9/2. Other centers are EPR invisible, *e.g.* [41]. A third S_3_ configuration with a broadened S_3_ signal was identified with ELDOR-detected NMR (EDNMR) in the presence of glycerol [42,43] and in PSII/Sr [44]. Although in [44] the authors did not completely rule out the presence of a closed cubane, five-coordinate S_3_ form, at the origin of this EPR signal, they favored a perturbation of the coordination environment at Mn4 and/or Mn3 in an open cubane S_3_ structure induced by glycerol. With X-and Q-band EPR experiments performed in the S_3_-state of plant PSII, both in the perpendicular and parallel modes, a high-spin, *S* = 6, was proposed to coexist with the *S* = 3 configuration. This *S* = 6 form was attributed to a form of S_3_ without O6/Ox bound and with the Mn^IV^_3_ part of the cluster in ferromagnetic interaction with the unsaturated dangler Mn^IV^ [43]. These two forms of S_3_ are, however, not detected by X-band EPR, so it seems unlikely that they correspond to the EPR invisible S_3_ mentioned above. Indeed, these new S_3_ signals described in [42,43] are detectable in the presence of glycerol and methanol, whereas the formation of the (S_2_ Tyr_z_^●^)′ state upon a near-IR illumination in the centers in S_3_ defined as EPR invisible is inhibited in the presence of glycerol (and in the presence of methanol in plant PSII) [45,46].

In cyanobacterial PSII/Sr [44], a proportion of centers exhibited a pulsed W-band field-swept S_3_ spectrum much broader than in PSII/Ca. This signal was proposed to be present in centers containing a 5-coordinate Mn ion in centers in which no water binding event takes place during the S_2_ to S_3_ transition. It was therefore proposed that, in these centers, the oxidation event would precede the water binding. Computational analyses also suggested heterogeneities in S_3_ [47–49] with also a *S* = 6 spin value [49].

None of the heterogeneities described above were detected in the crystallographic structures of S_2_ and S_3_ known to date (see references above). In addition, recent high-energy resolution fluorescence detected X-ray Absorption Spectroscopy together with QM calculations ruled out the presence of either peroxo or oxo/oxyl level intermediate to explain the S_3_ heterogeneity [50].

It is quite possible that some of the structural differences that cause the differences identified in EPR are too small to be detectable given the resolution of the crystallographic data. This at least shows, if it were necessary, that spectroscopy remains an indispensable complement to crystallography.

The EPR data summarized above describes a static view of the trapped configurations. Kinetically, it is well documented that the transition from S_2_ to S_3_ involved at least two phases. The fastest phase, with a *t*_1/2_ ≤ 25 μs, is attributed to a proton transfer/release. This fast phase precedes the electron transfer from S_2_ to the Tyr_z_^●^ occurring with a *t*_1/2_ ≤ 300 μs [51–55], and the binding of O6/Ox to the Mn1, *e.g.* [56]. It was proposed that the fast phase could correspond to the release of a proton in an intermediate step S_2_^LS^Tyr_z_^●^ → S_2_^HS^Tyr_z_^●^ before the S_2_^HS^Tyr_z_^●^ → S_3_*^S^*^=3^ transition occurs [32]. The existence of intermediate states in the S_2_ to S_3_ transition was tracked by following, at room temperature, the structural changes in the S_2_ to S_3_ transition in the µs to ms time-range after the 2nd flash [17,56]. No indication was found for a closed cubane intermediate. However, there was no conclusion on the spin state of the intermediate forms of S_2_ able to progress to S_3_. By definition, an intermediate state has a low concentration that makes its detection difficult and thus, the question of the existence of a high spin intermediate state in the S_2_ to S_3_ transition remains. It is possible that the S_2_^HS^ corresponds to a subtle structural/tautomeric intermediate that nonetheless mediates proton release during the S_2_^LS^Tyr_z_^●^ → S_2_^HS^Tyr_z_^●^ as a prerequisite to the formation of the S_3_ *S* = 3 state. It has been suggested that the fast phase observed in this transition corresponds to a proton release/movement associated with the formation of a S_2_^HS^ state [32,37]. If this is correct, we would expect to detect a change in the flash pattern of the proton release in conditions in which the S_2_^HS^ EPR signal is the flash-induced state. This was indeed the case since we have kinetically detected a proton release in PSII/Ca and PSII/Sr at pH 6.0 and 7.0, knowing that at pH 7.0, in PSII/Sr in contrast to PSII/Ca, half of the centers exhibit the S_2_^HS^ signal at g ∼ 4.8 able to progress to S_3_ at low temperatures [37].

Independently of the pH effect and Ca/Sr exchange, several mutations, the list being too long for making it here, in the first and second coordination sphere of the Mn_3_O_4_Ca unit strongly perturb the water splitting process. One of these mutants is the D1/V185N mutant in *Synechocystis* sp. PCC 6803 [57–60]. The D1/V185 residue is located adjacent to the cluster near three potential substrate candidates (W2, W3 and O5). It may also block or regulate the access to the open coordination site at Mn1. The D1/V185N mutation in *Synechocystis* sp. PCC 6803 slows down the release of O_2_ from 1.2 ms to about 27 ms halftime at 27 °C. In addition, the rate of exchange for the slower exchangeable water substrate molecule, Ws, was increased in the S_1_ and S_2_ states in the D1/V185N mutant, while both Wf, the faster exchangeable water substrate molecule, and Ws exchange rates were decreased in the S_3_ state in this mutant [60]. In contrast to the situation in the V185T mutant in *Synechocystis* 6803 in which the oxygen release kinetics was hardly affected (from 1.2 ms to 1.5 ms) [58], the D1/V185T mutation in *T. vestitus* resulted in similar phenotype to those in the D1/V185N in *Synechocystis* sp. PCC 6803 [61].

In the D1-V185T mutant in *T. vestitus* we found that the S_2_ state was mostly present in a high spin configuration [61] although a modified multiline was also observed. This observation triggered us to probe the proton release in the S_1_ to S_2_ transition in the context of the model mentioned above where the proton released in the S_2_ to S_3_ transition occurs in the S_1_Tyr_z_^●^ → S_2_^HS^Tyr_z_^●^ transition prior to the S_2_^HS^Tyr_z_^●^ to S_3_ transition [32,37]. One of the challenges here is that measuring proton release kinetics using a dye, as we did in [61], requires extensive washing of the PSII samples to remove all buffers. This treatment can have deleterious effects on the particularly fragile D1/V185T mutant. In so doing, we observed a rapid phase of proton release into the bulk, the kinetics of which was close, and difficult to distinguish from that observed following oxidation of Tyr_Z_ at pH 6.3 in Mn-depleted PSII. To get around this problem, in the present work, we also measured the electrochromic band shifts induced by each of the flashes in a sequence that is an alternative way to follow the movement of the charges in and around the Mn_4_CaO_5_ cluster [37,55].

As the μs components kinetics in the P_680_^+^ reduction reflect the progressive shift to the left of the equilibrium P_680_^+^Tyr_Z_ ↔ P_680_Tyr_z_^●^ resulting from slower proton release/movements [62–65], we also measured the P_680_^+^ decay in the D1/V185T mutant in *T. vestitus*.

Recently [60], it was claimed that the high spin EPR signal in the S_2_ state of the D1/V185N PSII in *Synechocystis* 6803 was not detectable. We report here, in an independent EPR experiment, that the S_2_^HS^ EPR signal in the D1/V185N PSII from *Synechocystis* 6803 is clearly present. This combined with the retardation in the S_n_ state transition reaction kinetics, now provides a consistent picture of the effects mutations at this second sphere location and reinforces criticality of the H-bonding network in in mediating the catalysis of water oxidation.

## Materials and methods

Purification of the D1-V185N mutant from *Synechocystis 6803* was previously described in [57]. The D1 protein is PsbA2 in this mutant strain. Purification of the D1/V185T mutant from *T. vestitus* was previously described in [61]. The D1/V185T mutant has been constructed in the *psbA_3_* gene so that the wild type PSII is that one with PsbA3.

Time-resolved absorption changes measurements were performed with a lab-built spectrophotometer [66] slightly modified as described in detail in [67]. For the ΔI/I measurements the samples were diluted in 1 M betaine, 15 mM CaCl_2_, 15 mM MgCl_2_, and 40 mM MES (pH 6.5). PSII samples were dark-adapted for ∼1 h at room temperature (20–22°C) before the addition of 0.1 mM phenyl *p*–benzoquinone (PPBQ) dissolved in dimethyl sulfoxide. The chlorophyll concentration of all the samples during the measurements was ∼ 25 μg of Chl mL^−1^. After the measurements, the ΔI/I values were normalized to exactly *A*673 = 1.75, that is very close to 25 μg Chl mL^−1^. The measurements with the dye bromocresol purple were done, after the removal of Mes, as previously reported [37].

X-band cw-EPR spectra were recorded with a Bruker Elexsys 500 X-band spectrometer equipped with a standard ER 4102 (Bruker) X-band resonator, a Bruker teslameter, an Oxford Instruments cryostat (ESR 900) and an Oxford ITC504 temperature controller. The samples, at ∼ 1.35 mg Chl mL^−1^, were dark-adapted for 1 h on ice then the samples were frozen in the dark (without any addition) to 198 K in a dry-ice/ethanol bath and then transferred into liquid N_2_ (77 K). Prior to recording the spectra, the samples were degassed at 198 K. The S_2_ state was induced by illumination with an 800 W tungsten lamp filtered by water and infrared cut-off filters for approximately 5–10 s at 198 K in a nonLsilvered Dewar in ethanol cooled with dry ice.

## Results

As the interpretation of the data strongly depends on the presence of an S_2_^HS^ state in the D1/V185N mutant in *Synechocystis* 6803, something that was contested in [60], we will firstly show the EPR spectra recorded in this strain. Fig. 1 compares the spectra in the PSII from *Synechocystis* 6803 WT (Panel A), and from the *S* 6803 D1/V185N-PSII (Panel B). The black spectra were recorded in the dark-adapted samples, the red spectra after continuous illumination at 198 K for 5-10 s, and the blue spectra are the light-*minus*-dark difference spectra.

**Figure 1:**
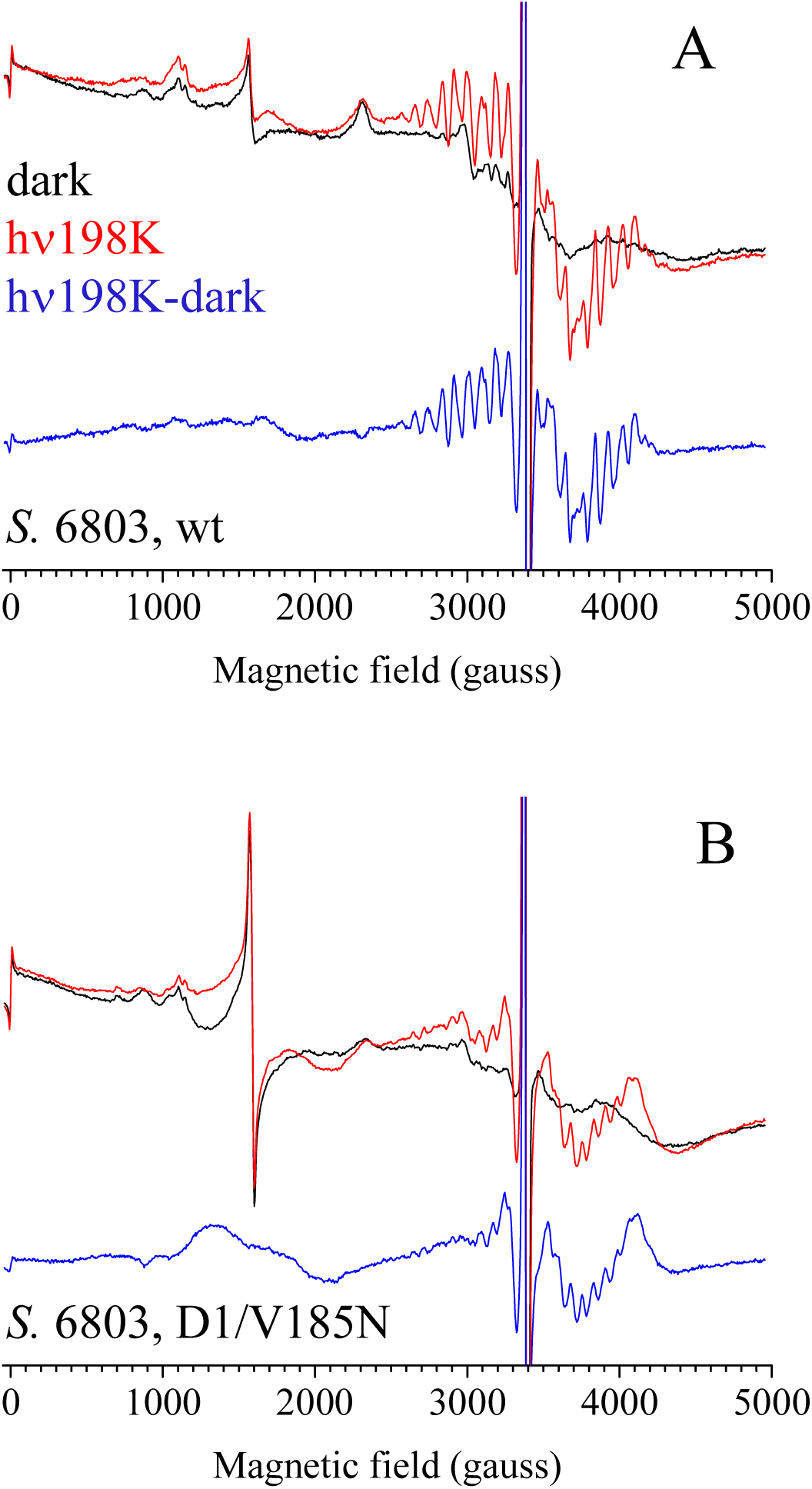
EPR spectra recorded at 8.6 K in WT PSII from *Synechocystis* 6803 (Panel A) and D1/V185N PSII from *Synechocystis* 6803 (Panel B). The black spectra were recorded in dark-adapted PSII. The red spectra were recorded after an illumination at 198 k for 5-10 s. The blue spectra are the “light”-*minus*-“dark” difference spectra. Instrument settings: [Chl] ∼ 1.35 mg /ml; modulation amplitude, 25 G; microwave power, 20 mW; microwave frequency, ∼ 9.49 GHz; modulation frequency, 100 kHz. The unresolved spectral region at g ∼ 2 (∼ 3390 gauss) corresponds to the saturated signal from Tyr_D_^●^.

In the WT PSII (Panel A) the light-induced state exhibits a normal S_2_^LS^ multiline signal similar to that induced in plant PSII and cyanobacterial PSII [10,24–26,30,32]. In contrast, in the D1/V185N PSII (Panel B), the shape and the amplitude of the multiline signal differs significantly from that one in the WT PSII. The S_2_ multiline signal in the D1/V185N sample is very similar, if not identical, to the S_2_ multiline signal in the D1/V185T mutant in *T. vestitus* [61]. Importantly, a large S_2_^HS^ EPR signal, centered at ∼ 1700 gauss, was induced by the illumination at 198 K in the D1/V185N mutant. This S_2_^HS^ EPR signal is also virtually identical to the S_2_^HS^ EPR signal detected in the D1/V185T mutant in *T. vestitus* [61].

In the black and red spectra other signals are detected. The *g*_z_ and *g*_y_ of Cyt*c*_550_ are detected at ∼ 2300 and ∼ 3000 gauss, respectively. The two small negative signals in the blue spectra, between ∼ 800 and ∼ 1000 gauss, correspond to the negative non-heme iron signal. This means that the non-heme iron was oxidized in a proportion of centers in the dark-adapted state and reduced by the Q_A_^−^ formed by the illumination at 198 K [68]. The large narrow signal at around 1600 gauss is due to contaminating Fe^3+^. In both samples, the *g* ∼ 1.9 form of the Q_A_^−^Fe^2+^Q_B_ signal (∼ 3700 gauss) and the Q_A_^−^Fe^2+^Q_B_^−^ biradical signal at *g* ∼ 1.6 (∼ 4400 gauss) are detected. The change in their relative proportion is likely due to a different proportion of the Q_A_Fe^2+^Q_B_^−^ state in the dark-adapted samples. In the black spectra, the Q_A_Fe^2+^Q_B_^−^ signal is possibly present at ∼ 4200 gauss and indeed larger in the mutant, *i.e.* [69].

In conclusion, an unequivocally similar S_2_^HS^ EPR signal is formed in the D1/V185N PSII from *Synechocystis* 6803 and the D1/V185T PSII mutant from *T. vestitus* mutant [61].

Knowing with certainty that both mutants exhibit an S_2_^HS^ state at pH 6.5, the question about a proton release into the bulk in the S_1_ to an S_2_ state with such S_2_^HS^ EPR properties had to be addressed again. Firstly, by measuring the proton/release using the dye bromocresol purple. Secondly, by recording the electrochromic band-shifts in the Soret region of the P_D1_ absorption spectrum at 440 nm. Indeed, this measurement takes into account both the proton uptake/release and the electron transfer events, *e.g.* [55,70] and references therein. In addition, unlike the situation with the dyes, the electrochromism is not contaminated by the protonation/deprotonation events coupled to the reduction/oxidation of the non-heme iron, and its measurement does not require an extensive washing of the PSII sample. For the removal of the contributions due to the reduction of Q_A_ the ΔI/I at 424 nm was also measured. Indeed, the electrochromism due to Q_A_ ^−^ equally contributes at 440 nm and 424 nm [70]. These measurements can be compared to the proton uptake/release kinetics again measured with bromocresol purple as in [61].

Panels A and B in Fig. 2 shows the absorbance changes of bromocresol purple in PsbA3-PSII and PsbA3/V185T-PSII, respectively, both from *T. vestitus*. An uptake of protons by PSII induces an increase in the ΔI/I at 575 nm, and a release of protons into the bulk by PSII induces a decrease of the ΔI/I. After the first flash (black points), there was a large increase in the absorption in both samples. This increase is due to the proton uptake following the reduction of the non-heme iron [37]. In PsbA3-PSII (Panel A) there is no detectable proton release associated with the S_1_ to S_2_ transition as already seen in such a PsbA3-PSII [37]. This is consistent with a large body of experiments indicating the absence of proton release in this transition, and with an approximately 1-0-1-2 pattern of proton release for the S_0_ to S_1_, S_1_ to S_2_, S_2_ to S_3_ and S_3_ to S_0_ transitions, reviewed in [37,57]. In contrast with our previous study in the V185T mutant from *T. vestitus* [61], the amplitude of the decrease in the ΔI/I in the 10 µs to 100 µs time range is very small, suggesting that a proton release, if any, occurs in a very low proportion of the centers. This lack of proton release is very likely due to the use of a PsbA3/V185T-PSII sample less damaged by the washing steps compared to the earlier study. Indeed, the much less damaged PSII core samples is also evident in the retention of stronger period four oscillations evident in the measurements of flash-induced absorption changes measured at 432 nm, as shown in Figure 4, Panel E, in comparison to the earlier work [61] as further discussed below. Additionally, proton uptake is detected after each flash as previously reported [37] explaining the drift in the signal (not subtracted in Panels A and B) observed on each flash. The proton release kinetics in the PsbA3/V185T-PSII in the S_2_ to S_3_ transition (red points) and in the S_0_ to S_1_ transition (green points) appears hardly affected when compared to PsbA3-PSII (∼ 30-40 µs, and ∼ 100-200 µs, respectively). In the S_3_ to S_0_ transition, the release of the first proton is also hardly affected in the mutant. In contrast, the release of the second proton is strongly slowed down with a kinetics similar to the O_2_ release (∼ 20-30 ms) [57,61].

**Figure 2:**
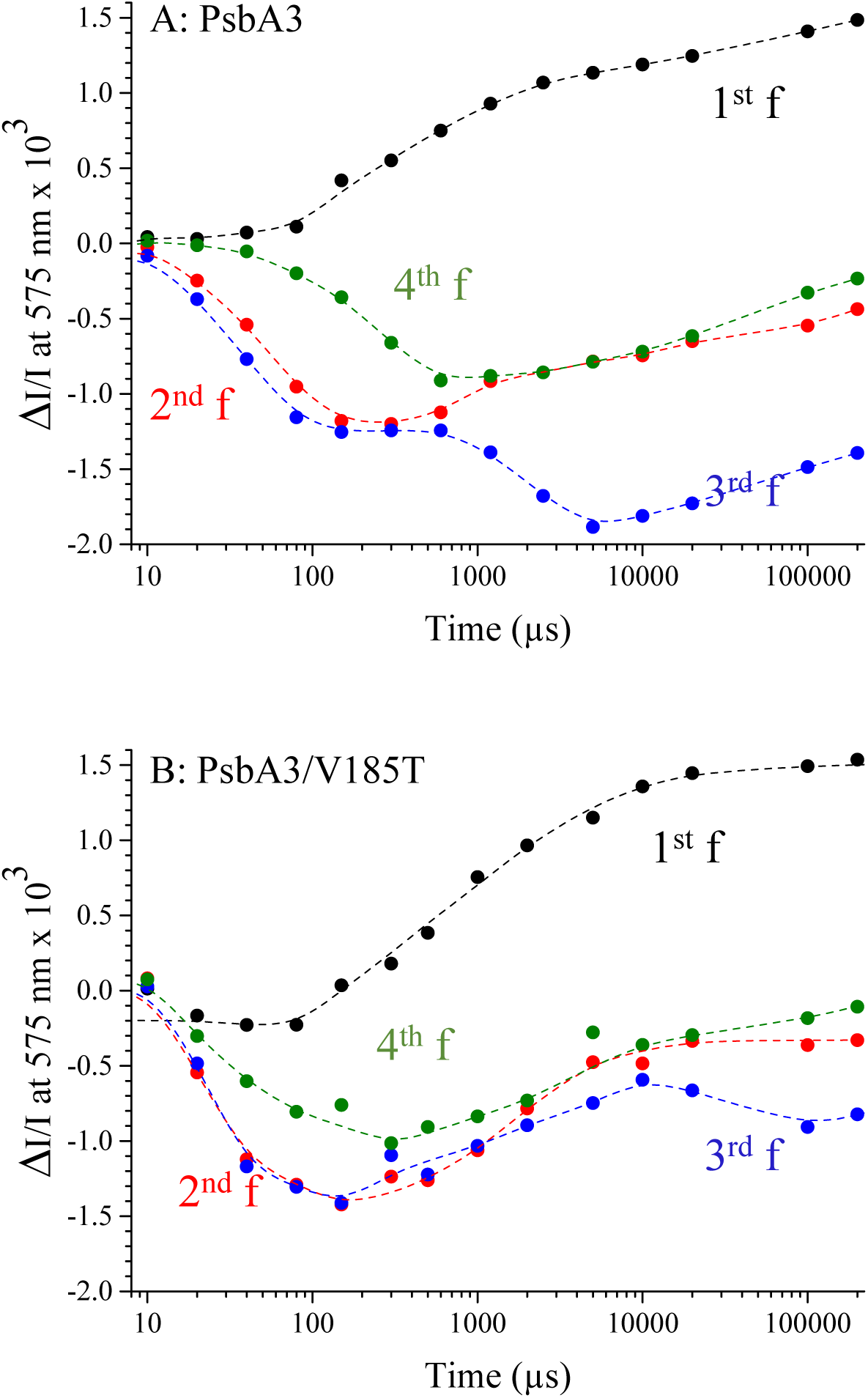
Time-courses of the absorption changes of bromocresol purple at 575 nm in PsbA3-PSII (Panel A) and PsbA3/V185T-PSII (Panel B) from *T. vestitus* after the 1^st^ flash (black points), the 2^nd^ flash (red points), the 3^rd^ flash (blue points), and the 4^th^ flash (green points). PPBQ (100 µM final concentration) dissolved in dimethyl sulfoxide was added after the dark-adaptation. Flashes spaced 400 ms apart. The dashed lines have been drawn to join the points.

**Figure 3:**
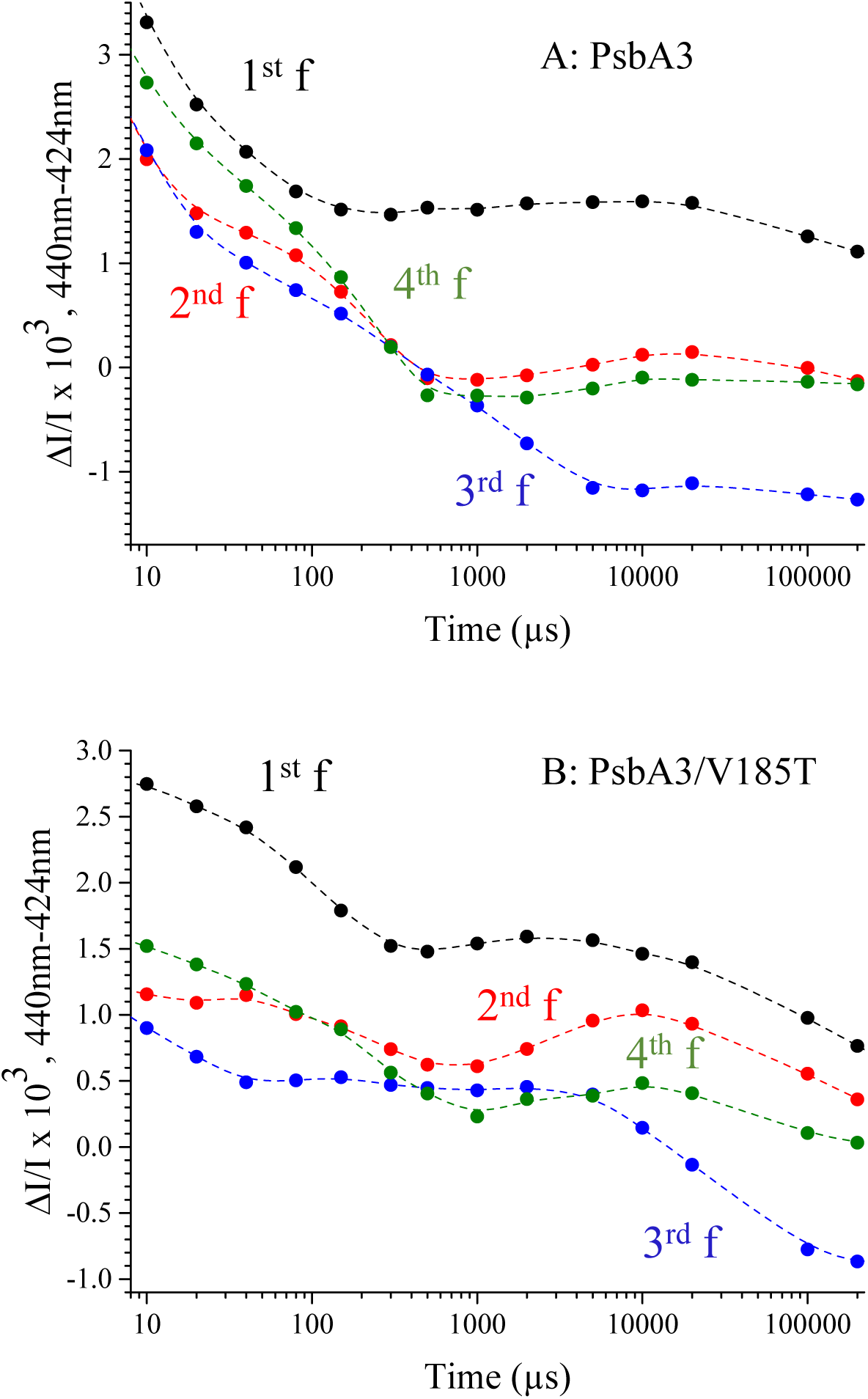
Time-courses of the absorption change differences 440 nm-*minus*-424 nm after the first flash (black), the second flash (red), the third flash (blue), and the fourth flash (green) given to dark-adapted PsbA3-PSII (Panel A) and PsbA3/V185T-PSII (Panel B) in the presence of 100 μM PPBQ with flashes spaced 400 ms apart. The dashed lines have been drawn to join the points.

**Figure 4:**
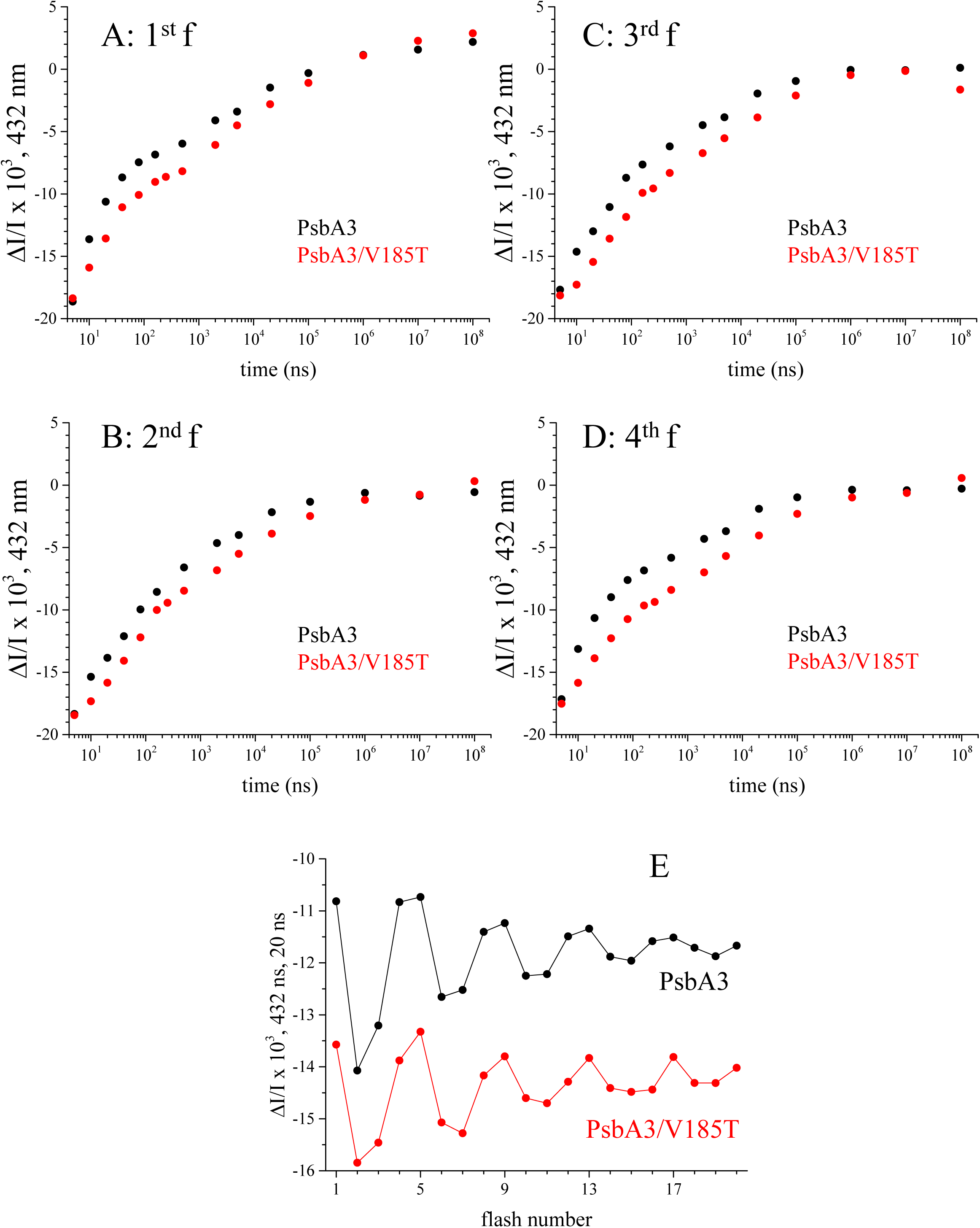
Kinetics of flash-induced absorption changes measured at 432 nm in PsbA3-PSII (black), and PsbA3/V185T-PSII (red) from *T. vestitus* after the 1^st^ flash (Panel A), the 2^nd^ flash (Panel B), the 3^rd^ flash (Panel C) and the 4^th^ flash (Panel D). The amplitudes of the kinetics were normalized to OD673nm = 1.75 (a Chl concentration of ∼ 25 μg/ml). 100 μM PPBQ was added before the measurements were taken. Panel E shows the amplitude of the ΔI/I at 432nm, 20 ns after each flash of the sequence with flashes spaced 400 ms apart.

Fig. 3 shows the kinetics of the 440 nm-*minus*-424 nm difference in PsbA3-PSII (Panel A), and PsbA3/V185T-PSII (Panel B) from *T. vestitus*. After the first flash (black points), the kinetics that corresponds to the S_1_Tyr_z_^●^ to the S_2_Tyr_z_ transition is slightly slowed down from 10-20 µs in PsbA3-PSII to 50-60 µs in the PsbA3/V185T mutant. After the second flash (red points) the kinetics is biphasic with a fast phase (*t*_1/2_ ∼ 10-15 µs) that we have attributed to a proton movement coupled to the S_2_^LS^Tyr_Z_^●^ to S_2_^HS^Tyr_Z_^●^ transition, and the slow phase was attributed to the S_2_^HS^Tyr_Z_^●^ to the S_3_Tyr_Z_ transition [37,55]. In the PsbA3/V185T-PSII mutant the two phases are not really differentiated, which makes the interpretation difficult even if this result could suggest a slowing down of the proton release in the S_2_ to S_3_ transition. In the PsbA3-PSII, after the fourth flash, the kinetics is also biphasic with the fast phase corresponding to the electron transfer in the S_0_ to S_1_ transition (*t*_1/2_ ∼ 10-15 µs), which precedes the proton release with a *t*_1/2_ ∼ 200 µs [37,51,55]. In the PsbA3/V185T-PSII, after the fourth flash (green points), although the two phases are also difficult to be detected due to a lower signal-to-noise ratio, the two phases seem to have more or less the same kinetics. The important difference between Panel A and Panel B is the slow phase after the third flash (blue points) with a *t*_1/2_ ∼ 20-30 ms in PsbA3/V185T-PSII instead ∼ 2 ms in PsbA3-PSII. The *t*_1/2_ of the slow phase (both in PsbA3-PSII and PsbA3/V185T-PSII) is similar to that of the proton release in these samples. After the 4^th^ flash the slow decay certainly originates from a residual S_3_ to S_0_ transition due to the misses but we have no clear explanation for the slow drift after the 1^st^ and 2^nd^ flash since the redox events at the non-heme iron do not contribute in the electrochromism of P_D1_.

Changes in the kinetics of electron and proton motions, should affect the Tyr_Z_P_680_^+^ ↔ Tyr_z_^●^P_680_ equilibrium and therefore the P_680_^+^ decay kinetics. Panels A, B, C, and D in Fig. 4 show the P_680_^+^ reduction kinetics measured at 432 nm in PsbA3-PSII (black), and PsbA3/V185T-PSII (red) after the 1^st^ flash (Panel A), the 2^nd^ flash (Panel B), the 3^rd^ flash (Panel C) and the 4^th^ flash (Panel D). In all the S_n_ states, the decay of P_680_^+^ in the PsbA3/V185T-PSII mutant was significantly slower both in the ns time-domain corresponding to the pure electron transfer step, and the µs time-domain where the electron transfer(s) is(are) coupled to proton movements. The Tyr_Z_P_680_^+^↔ Tyr_Z_^●^P_680_ equilibrium is therefore shifted to the left in the PsbA3/V185T-PSII mutant when compared to the PsbA3-PSII in all the S_n_ states [62,64] and we expect that some of the proton movements coupled to the electron transfer are also slowed down.

Panel E in Fig. 4 shows the ΔI/I measured 20 ns after each flash of the sequence in the PsbA3-PSII (black) and in the PsbA3/V185T-PSII (red). Due to the slower decay of P_680_^+^ in the S_2_ and S_3_ states [71] than in the S_0_ and S_1_ states we obtain an oscillation with the period of four. Since in all the S_n_ states the decay of P_680_^+^ is faster in PsbA3-PSII than in PsbA3/V185T-SII, the ΔI/I values in PsbA3-PSII, 20 ns after the flashes, are smaller (less negative) in all the S_n_ states. However, the oscillations are similar in both samples, which indicates that in the PsbA3/V185T-PSII used here the miss parameter is not significantly larger than in the PsbA3-PSII.

## Discussion

One of the main results in this work is the presence of a S_2_^HS^ in the D1/V185N-PSII from *Synechocystis* 6803, with EPR characteristics strikingly similar to those observed in the D1/V185T-PSII mutant from *T. vestitus* [61]. Although the S_2_^HS^ EPR signal in [60] is clearly recognizable, baseline distortions in the low-field region of the wild type spectrum appear to have prevented the authors from identifying the S_2_^HS^ EPR signal in the mutant. Therefore, these data reinforce the idea that the D1/V185 residue influences the high-spin/low-spin equilibrium of the S_2_ state in a similar manner across different cyanobacterial species and supports a consistent mechanism by which a perturbation of this site modulates the still enigmatic S_2_^HS^ state in PSII.

At first glance, it is surprising that the D1/V185T mutation in *Synechocystis* 6803 had almost no effect [57–59], in contrast to *T. vestitus*, where it induced the same phenotype as the D1/V185N mutation in *Synechocystis* 6803. However, studies of Photosystem II properties in *T. vestitus* with PsbA1, PsbA2, or PsbA3 as the D1 protein, we have frequently observed that site-directed mutations in PsbA1, PsbA2, or PsbA3 done in order to reproduce the situation in another PsbA did not fully result in the expected phenotype. For instance, the PsbA3/E130Q mutation did not yield a PSII with properties similar to those of PsbA1-PSII [72]. We attributed this to partial compensation by the approximately thirty additional amino acid differences associated with the distinct PsbA variants. It is plausible that the same phenomenon occurs here when comparing the V185T mutation in the PsbA3 D1 protein of *T. vestitus* with the same mutation in the PsbA2 D1 protein of *Synechocystis* 6803. Moreover, these sequence differences likely reflect broader evolutionary adaptations, as *Synechocystis* is a mesophile, whereas *T. vestitus* is a thermophile. Thermophilic proteins are characterized by a stiffer framework due to a higher density of non-covalent interactions, which enhances stability at high temperatures but increases the activation energy required for conformational changes [73]. In *T. vestitus*, this rigidity may amplify the D1/V185T mutation’s impact on the hydrogen-bond network around the Mn_4_CaO_5_ cluster, impeding the water dynamics of the formation of optimal H-bond network and proton transfer pathways at room temperature [57,74]. In contrast, the more flexible framework of the mesophilic *Synechocystis* enzyme may buffer the mutation’s effects, reducing its influence on proton-coupled electron transfer and substrate water stabilization. This interplay between protein rigidity, hydrogen-bond strength, and activation energy may therefore explain the differential impact of the D1/V185T mutation in the two organisms.

It has been shown that the equilibrium between the S_2_^LS^ and S_2_^HS^ states is pH dependent [32], with a pK_a_ ∼ 8.3 for the native Mn_4_CaO_5_ cluster and a significantly shifted pK_a_ of ∼ 7.0 for the Mn□SrO□ cluster [37]. Alkaline conditions are thought to mimic the charged state of the Tyr_z_^●●^, emulating the electrostatic proton ejection regime that facilitates progression through the Kok cycle [32,37,51,52,71,74–76]. Furthermore, there is some evidence that the Mn_4_CaO_5_ must pass through an S_2_^HS^ intermediate state before forming the S_3_ state. For example, the S_2_^HS^ state formed by the room temperature flash illumination could progress to S_3_ under further continuous illumination at 198 K. Similarly, samples advanced to the S□^LS^ state upon illumination at 198 K subsequently form the S_2_^HS^ state when warmed to room temperature, enabling progression to the S_3_ state under further illumination at 198 K [77]. These observations have led to the tentative conclusion that the S_2_^HS^ state is a necessary intermediate during the S_2_ → S_3_ transition.

The lowered pK_a_ with the Mn□SrO□ cluster explains why the S_2_^HS^ to S□ transition can occur in a significant proportion of centers even at pH 6.5. Interestingly, D1/V185N-PSII in *Synechocystis* 6803 and D1/V185T-PSII in *T. vestitus* exhibits a high proportion of centers with an S_2_^HS^ signal at pH 6.5, suggesting that these centers could progress to the S_3_ state under illumination at 198 K. However, this progression is not at all evident in D1/V185T-PSII with a Mn_4_CaO_5_ cluster, as shown in spectrum *a* of Fig. 9 in [61]. In contrast, in D1/V185T-PSII with a Mn_4_SrO_5_ cluster, the effect of the 198 K illumination was more pronounced (spectrum *b* in Fig. 9 in [61]). However, the S_2_^HS^ EPR signal that disappeared in D1/V185T-PSII with a Mn_4_SrO_5_ cluster upon the 198 K illumination had a shape similar to that in the wild type PSII (with both Ca and Sr in the cluster). This observation suggests some heterogeneity in the EPR properties of the S_2_^HS^ state in D1/V185T-PSII with a Mn_4_SrO_5_ cluster. Importantly, since the illumination at 198 K results in the formation of the S_2_^HS^ and S_2_^LS^ states in both species, it seems likely that each of these two states arises from two structurally different S_1_ states.

In the present work, we observed that the S_2_^HS^ state in D1/V185T-PSII exhibits EPR characteristics distinct from those of the wild type, alongside altered kinetics in proton release. The previously mentioned possibility [61] for a S_2_^HS^ to S_3_ transition occurring at 198 K in D1/V185T-PSII seemed to be supported by the detection of a proton release on the first flash. However, it is shown here that this proton release does not occurs in active D1/V185T-PSII but rather as originating from the oxidation of Tyr_Z_ in a minor sub-population of D1/V185T-PSII in which the washing procedure resulted a Mn-depletion. It can now be concluded that the S_2_^HS^ state in D1/V185T-PSII, with EPR properties different from those in wild type PSII, is unable to progress to S_3_ at 198 K, most likely because its protonation state and/H-bonding configuration differs. This is consistent with a model wherein the mutation introduces perturbations to the hydrogen-bonding network near the Mn_4_CaO_5_ cluster, resulting in a suboptimal protonation environment for progression to the S_3_ state. The lack of proton release during the S_1_ to S_2_^HS^ transition further supports this interpretation and aligns with earlier studies indicating that proton release is a prerequisite for the S_2_^HS^ to S_3_ transition in wild-type PSII.

The S_2_^HS^ EPR signal in the D1/V185T-PSII from *T. vestitus*, and the D1/V185N-PSII from *Synechocystis* 6803, have a shape reminiscent of the spectrum induced by near-infrared illumination of the S_2_^HS^ state at high pH in PsbA3-PSII see Fig. 3 in [77]. Presently, there is no explanation of the near-infrared illumination however this similarity could suggest that the two states only differ by small structural changes, such as shifts in the H-bonding and proton tautomer configurations.

Functionally significant hydrogen bonding changes due to mutations at D1/V185 likely affect the stabilization and dynamics of substrate water molecules in the S_3_ state based upon previous findings [57,59,60]. The slowdowns of both the lag and oxygen release phase during the S_3_ to S_0_ transition have been attributed to changes in the H-bond network that correspond to a suboptimal configuration of reactants and an associated entropic penalty for crossing the activation barrier of the transition state [57]. The observation of slower water exchange in S_3_ [60] has been suggested to be the result of the stabilization of water molecules due to altered H-bonding around the inserted substrate water, O_6_/O_x_. However, the mechanism of substrate exchange remains to be elucidated. As noted, D1/V185T and D1/V185N mutations replace the non-polar valine side chain with threonine or asparagine, which can introduce non-native hydrogen bonds to the putative O_6_/O_x_ substrate and with water molecules (*e.g.* W3). This disruption likely affects both water binding and proton release.

The findings presented here underscore the central role of the H-bond network in modulating the LS-to-HS transition and subsequent steps in the Kok cycle. In Ca-PSII, the LS-to-HS transition is characterized by cooperative interactions among multiple protonatable groups, as evidenced by the best-fit value of n ∼ 4.4 [32], although only one proton is ultimately released to the bulk solution [37]. Here, we can speculate that this cooperativity lowers the transition state barrier for proton release rather than stabilizing the deprotonated HS state itself. According to this view, the well-organized H-bond network facilitates a collective reorganization that creates an energetically favorable pathway for the LS-to-HS transition, ensuring efficient coupling of proton dynamics with substrate water insertion at the Mn_4_CaO_5_ cluster.

This may provide a view of some shared characteristics of proton transfer during both the S_2_ → S_3_ and S_3_ → S_0_ transitions. This S_2_ → S_3_ transition also coincides with the critical insertion of W3, initially coordinated to Ca, into the open coordination site on Mn1. This step is crucial for positioning W3 to become Ox, one of the two substrate waters for the water oxidation reaction. The timing and coordination of W3 insertion and its deprotonation is expected to be strongly influenced by the surrounding H-bond network. In the LS state, the network stabilizes W3 on Ca²□ in a protonated form, maintaining readiness for the subsequent catalytic steps. Upon insertion into Mn1 following the LS-to-HS transition, the reorganization of the H-bond network facilitates the deprotonation of W3, with the cooperative action lowering the activation barrier for this step and ensuring the system is primed for subsequent oxidation chemistry. Similarly, simulations in [74] emphasize that concerted proton shuffling between O6 and the waters W3 and W2, along with the extremely wide preorganization of an extended hydrogen bond network, is mandatory for Tyr_z_^●^ reduction in the S_3_ → S_0_ transition. While a similar proton cooperativity to that seen in the formation of the S_2_^HS^ state, such a cooperative transition on the H-bond network and its disruption by mutations or Sr-substitution could also account, for the slowdown of the S_3_ → S_0_ transitions and help explain the increase in the entropic penalty and H/D effects previously observed for V185N. This insight aligns with the idea that efficient catalysis requires not just local H-bond interactions but a globally integrated network capable of synchronizing proton and water dynamics across the system. Disruptions to this network, as seen in D1/V185 mutations and Sr²□ substitutions, likely hinder the necessary concerted movements, leading to slowed transitions and impaired water oxidation.

Consistent with this interpretation, we observe that the proton release during the S_3_ → S_0_ transition in the D1/V185 mutants was primarily affected by the second of the two protons released during that transition. In contrast, the first proton release, which occurs rapidly following the formation of the tyrosine radical, Tyr_z_^●^, remains largely unaffected. This initial fast proton release is proposed to be linked to the electrostatic impact of uncompensated charge formation upon the effects of the formation of Tyr_z_OH-His190 → Tyr_z_O^●^-H^+^His190 [74,75]. Functionally, this is proposed to occur to prepare Asp61D1 as a proton acceptor for the slower, second proton release from substrate water O6 [74–75,78]. The transient electric field during the Tyr_z_O^●^-H^+^His190 state is calculated to promote proton transfer from W1 to Asp61, inducing rearrangements in the cluster of residues D1-Asp61, D1-Glu65, and D2-Glu312 into a configuration that facilitates rapid proton egress to the bulk [74–75,78]. The rapid exit of this proton is likely gated, preventing back-reactions. Once deprotonated, the cluster forms the base that accepts the proton from substrate water O6, facilitating the final electron transfer leading to O_2_ formation [74]. The fact that the fast proton release remains relatively unaffected by the mutation, while the second proton release is delayed (similarly to the O_2_ evolution), is consistent with the D1/V185 residue being distant from the fast proton release vector but positioned near the substrate water (O6), where deprotonation occurs. Thus, the delayed second proton release in the mutants is attributed to the disruption of the hydrogen-bonding network around O6, which slows the proton transfer pathway necessary for its deprotonation and the final stages of water oxidation.

## Acknowledgements

This work has been in part supported by (i) the French Infrastructure for Integrated Structural Biology (FRISBI) ANR-10-INBS-05, (ii) the Labex Dynamo (ANR-11-LABX-0011-01), (iii) the JSPS-KAKENHI Grant in Scientific Research on Innovative Areas JP17H064351, JSPS-KAKENHI Grant 21H02447 and a grant from the US-National Science Foundation (MCB1716408). One of the PSII preparations used in this work was done by T. Tibiletti.

## Abbreviations

Photosystem II, PSII; Chl, chlorophyll; Chl_D1_/Chl_D2_, monomeric Chl on the D1 or D2 side, respectively; MES, 2-(*N*-morpholino) ethanesulfonic acid; P_680_, primary electron donor; P_D1_ and P_D2_, individual Chl on the D1 or D2 side, respectively, which constitute a pair of Chl with partially overlapping aromatic rings; Phe_D1_ and Phe_D2_, pheophytin on the D1 or D2 side, respectively; PPBQ, phenyl *p*–benzoquinone; Q_A_, primary quinone acceptor; Q_B_, secondary quinone acceptor; Tyr_Z_, the tyrosine 161 of D1 acting as the electron donor to P_680_; WT*3, *T. elongatus* mutant strain deleted *psbA_1_* and *psbA_2_* genes and with a His-tag on the carboxy terminus of CP43. EPR, Electron Paramagnetic Resonance. ML, multiline. QM/MM, quantum mechanics/molecular mechanics.

